# Inferring Protein Domain Semantic Roles Using word2vec

**DOI:** 10.1101/617647

**Authors:** Daniel Buchan, David Jones

## Abstract

In this paper, using word2vec, we demonstrate that proteins domains may have semantic “meaning” in the context of multi-domain proteins. Word2vec is a group of models which can be used to produce semantically meaningful embeddings of words or tokens in a vector space. In this work we treat multi-domain proteins as “sentences” where domain identifiers are tokens which may be considered as “words”. Using all Interpro (Finn, Attwood et al. 2017) eukaryotic proteins as a corpus of “sentences” we demonstrate that Word2vec creates functionally meaningful embeddings of protein domains. We additionally show how this can be applied to identifying the putative functional roles for Pfam (Finn, Coggill et al. 2016) Domains of Unknown Function.

## Introduction

Word2vec (Mikolov 2013) is a group of models which can be used to learn the embeddings of words in a continuous vector space given a corpus of sentences. Often Natural Language Processing (NLP) tasks consider words as sets of unrelated tokens, subjecting them to no-more rigorous analysis than frequency counting. While this is mathematically and computationally convenient it ignores the fact that most words have degrees of similarity, such as verbs with differing tenses, adverbs with differing endings or words which share the same suffixes. Word2vec aims to produce embeddings of words in a vector space where distance in the vector space correctly encodes the degree to which words or terms are similar or can be used in similar semantic context. Although a great degree has been written about these methods it remains unclear exactly why these models are performant (Goldberg 2014). Nevertheless they show good performance in the task of clustering words with related semantic meaning, interested readers should consult the following paper for further details (Mikolov 2013). Since lexical word embeddings have become popular, they have been adapted and applied directly to protein and gene sequences. prot2vec, gene2vec and seq2vec are examples of such methods (Asgari and Mofrad 2015, Yang, Wu et al. 2018).

Proteins are often composed of discrete domains. These are either conceptualised as subsequences of independent protein sequences which share homology (and by extension evolutionary origin) (Finn, Coggill et al. 2016). Or alternatively domains may be considered structurally, where they are subsections of the proteins which are compact, independently folding and observed to be shared between a variety of proteins (Andreeva, Howorth et al. 2014, Cheng, Schaeffer et al. 2014, Dawson, Lewis et al. 2017). An extension of the observation that proteins can be decomposed in to sets of domains is the hypothesis that domains act as sub-functional units and when composed together a protein’s given combination of domains is what gives rise to the protein’s specific function (Das and Orengo 2015, Nepomnyachiy, Ben-Tal et al. 2017). In the following study we show that protein domains can be embedded in a “semantically” meaningful vector space and that this embedding space reflects meaningful information about the functional roles (in terms of GO term assignments) of the protein domains.

In the following work we briefly discuss the use of Word2vec in protein domain functional inference. Protein function prediction has received a great deal of attention in the preceding 20 years (Friedberg 2006) and a great number of function prediction methods have been developed. Many of these make use of sequence search using some manner of nearest neighbour functional assignment (Watson, Laskowski et al. 2005, Loewenstein, Raimondo et al. 2009). As the field has progressed work has been done to integrate more sophisticated statistical methods and models with many contemporary methods leveraging machine learning with ensemble or meta-prediction methodologies. Current state of the art in protein function is measured by the Critical Assessment In Function Annotation (CAFA) community experiment (Radivojac, Clark et al. 2013). In this experiment groups attempt to predict experimentally validated Gene Ontology (GO) terms (Consortium 2017) over a blind set of unannotated protein sequences. Thus far, performance and progress in this task indicates that protein function prediction remains a challenging problem in the field of bioinformatics.

## Method

### Datasets

We downloaded Interpro 62 (Finn, Attwood et al. 2017) with the associated GO and protein domain assignments. The files were parsed to extract only the Eukaryotic proteins and their GO and Pfam protein assignments. The following work looks only at eukaryotic proteins as there are few multidomain proteins in the bacteria and archaeal kingdoms, as such little domain context information would be available for proteins from those kingdoms. Only GO assignments with the following evidence codes were retained: EXP, IBA, IDA, IEP, IGC, IGI, IMP and IPI. This eliminates all the high throughput and more tenuous computational annotation assignments. The resulting dataset contains 9,030,650 eukaryotic proteins, which have domain assignments 11,355 of the available Pfam domain families and these proteins are associated with annotations from 2,358 GO Terms.

Not all regions within each protein have been assigned to domains. In large part because not all domains are known and assigned but also because many Eukaryotic proteins possess regions of intrinsic disorder (Walsh, Giollo et al. 2015), regions of low complexity or coiled coiled sequences. All such unassigned regions were compiled (see below). As Word2vec analyses words based on the semantic context of neighbouring words representing unassigned regions in our corpus preserves important domain context information.

These data were then used to derive which Pfam domains are seen to be associated to which GO terms. For every Pfam domain we associated all GO terms assigned to all the proteins the Pfam domain was observed in. This assigns a varied bag of GO terms to each Pfam domain and this bag of terms can be viewed as representing the spectrum of observed functional diversity for that Pfam domain.

### Unassigned sequence region assignments

The fasta sequence database for Interpro 62 was masked using pfilt for both coiled coil and low complexity regions using pfilt (Jones 1999). Disordered regions were derived directly from the Interpro disorder annotations. Gap regions which did not contain disorder annotations, coiled-coil or low complexity sequence were assigned given the length of the unassigned regions. These remaining gap regions were binned into size bins based on their lengths (see figure 1). The majority of gap regions are around 100 residues in length, as the typical structural domain size is around 100 residues 5 gap types were created to represent unassigned regions of various sizes which are approximate multiples of the typical domain size, see table 1. All non-domain regions: gaps, disordered, low complexity and coiled-coil regions were then compiled as a set of adjunct domain-like sequence regions to complement the PFAM domain assignments.

**Figure 1:**
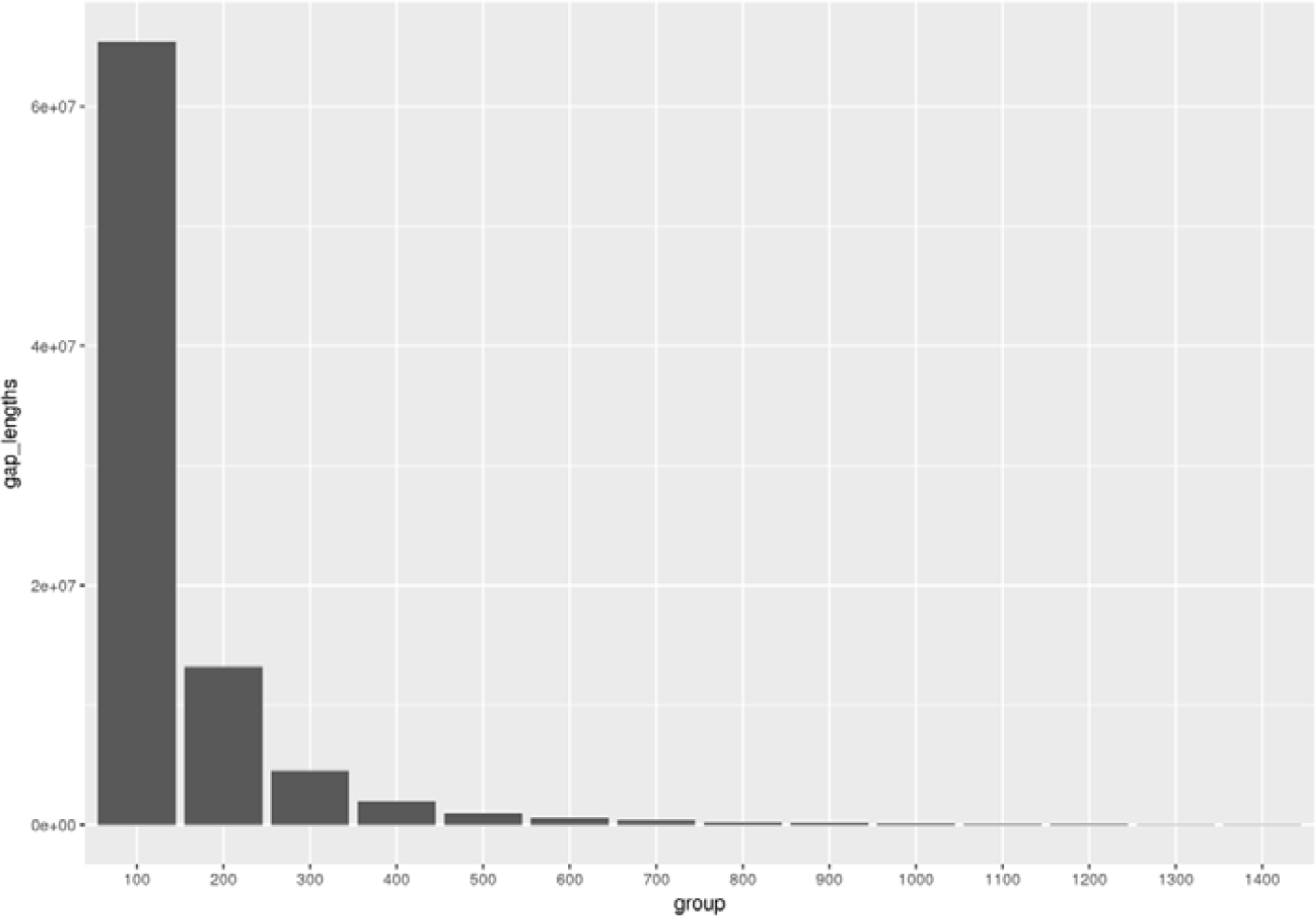
Distribution of gap regions (regions without Pfam domain assignments) in interpro Eukaryotic sequences.

**Table 1:** Names and sizes of gap pseudo-domains

| Gap Region ID | Size (residues) |
| --- | --- |
| G100 | 20-100 |
| G200 | 101-200 |
| G300 | 201-300 |
| G400 | 301-400 |
| G500 | 401- >500 |

### Building the word embedding

To build word2vec embeddings we treat protein sequences and their domain assignments as “sentences”. The Pfam IDs and other sequence region assignments are used as tokens/pseudo-words in such a pseudo-sentence. For instance a typical protein may be converted to a sentence such as “PF00170 PF003534 G200 LowComplexity PF00678”. Which would indicate two leading Pfam domains followed by a gap region up to 200 residues, a region of low complexity sequence finally terminating in a Pfam domain (see figure 2). We compile such sentences for every Eukaryotic protein in Interpro62 and this set of sentences becomes the corpus we use to create the word embedding.

**Figure 2:**
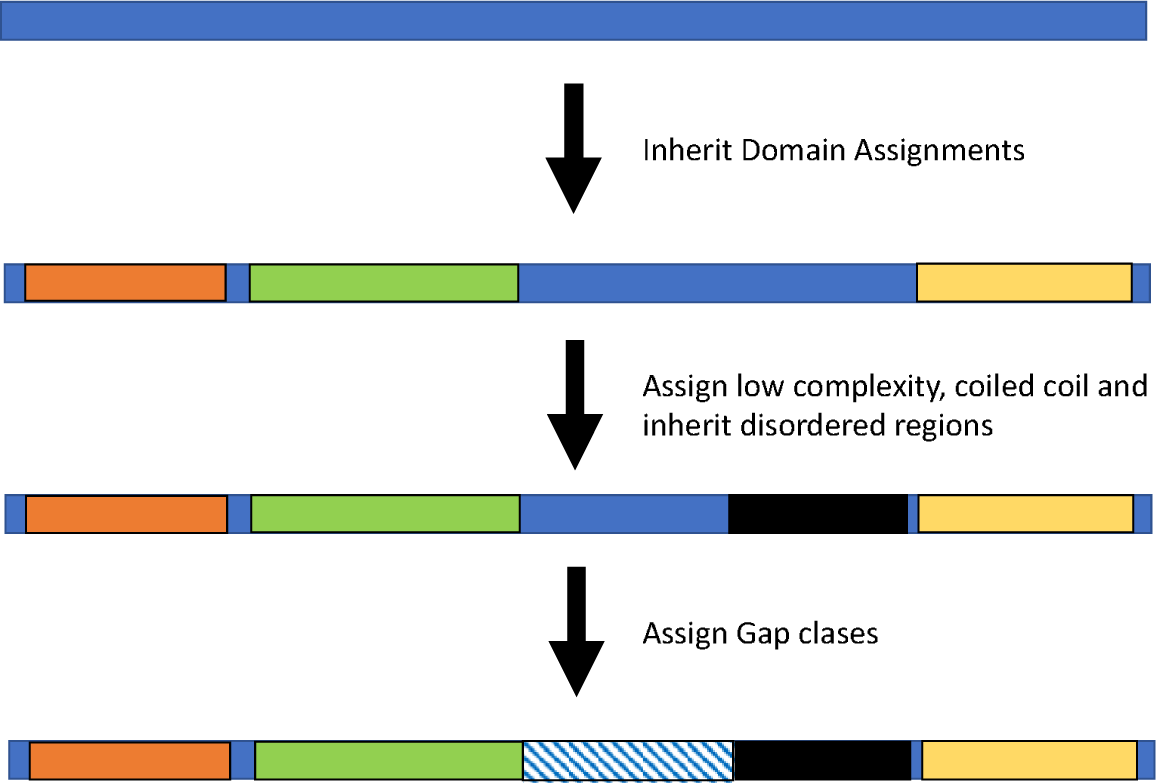
The example of the domain and saequence region assignement. Pfam domains and disorder regions are derived from Interpro annotations. Low Complexity and Coiled Coil regions are calculated by Pfilt and gaps are assigned given their size.

Python library genism (https://radimrehurek.com/gensim/) was used to create the word2vec model from the corpus using the default parameters. This means the embedding uses the skip-gram algorithm and model to build the embedding. This process is illustrated in full in figure 3. For the benchmark below an all-against-all distance matrix of domains was derived.

**Figure 3:**
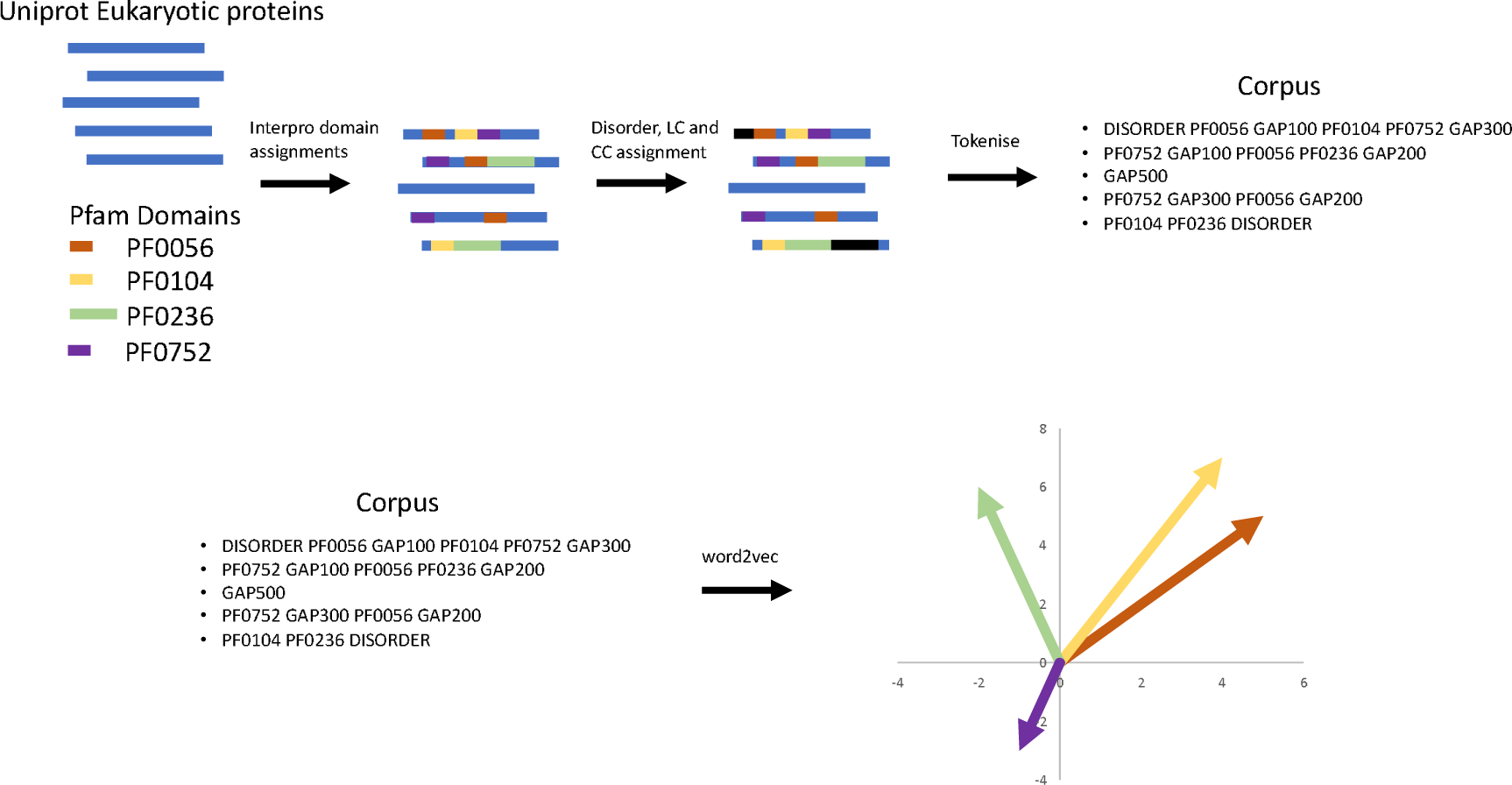
Compiling protein “sentences”. Interpro compiles assignments of domains on Uniprot protein sequences. We take only the Pfam domain assignments and complement those with assignments of Disorder, Low Complexity (LC) and coiled-coil (CC) regions. These are then tokenised to create a corpus of “sentences”. The corpus can then be used as input to word2vec. The output is a vector space which places each token at a point within that space, here stylised in 2D. Tokens which appear in similar syntactic contexts in the corpus should be placed near one another in the vector space.

### Benchmark

We are interested in whether word2vec embeds Pfam domains in a manner which is biologically meaningful. This would in turn would indicate that there is some manner of semantic meaning in the positioning or sequence context for protein domains. To investigate the embedding, initially we attempted to project the domain vectors into three dimensions (data not shown) using Multi Dimensional Scaling. However the resulting projection did not yield any trivially interpretable result.

An alternative means of investigating whether the embedding is biologically meaningful would be to establish if functionally related domains are placed near one another in the embedding. To investigate this we assigned GO terms to the Pfam domains. This was done by allowing Pfam domains to inherit all GO terms assigned to the proteins each Pfam domain is observed in. Although this is somewhat lossy, as GO annotations reflect protein functions, each domain’s “bag” of GO terms will reflect the functional diversity for the domain. 2,358 GO terms were assigned over the 11,355 Pfam domains observed in the Eukaryotic proteins. These assignment could then be used for a nearest neighbour benchmark test.

## Results

### Nearest Neighbour Performance

Performance in nearest neighbour functional annotation was calculated to assess whether the vector embedding of domains displayed any meaningful structure. That is, domains with similar functionality were placed near one another in the embedding. Each domain was in turn considered by inheriting the GO terms from its K nearest neighbours and comparing these predicted terms to the known terms assigned via Interpro annotations. Table 2 gives the precision and Mathew’s Correlation Coefficients (MCC) scores for the nearest neighbour benchmark. The MCC value indicates the predicted terms are non-random (greater than 0) which in turn suggests that there is some meaningful structure in the embedding of domains in a vector space.

**Table 2:** Mean precision and accuracy and Mathew’s Correlation Coefficients given nearest neighbour inheritance of GO terms.

| K Nearest Neighbours | Mean Precision | Mean Accuracy | Mean MCC |
| --- | --- | --- | --- |
| 1 | 0.33 | 0.99 | 0.28 |
| 3 | 0.19 | 0.98 | 0.23 |
| 5 | 0.15 | 0.98 | 0.22 |
| 10 | 0.10 | 0.96 | 0.19 |

Word2vec is trained to embed human language words in a vector space such that words which occur in similar semantic contexts are close to one another in the vector space. That our domain embedding is non-random implies that multidomain proteins exhibit some form of semantic structure. That is, certain domains appear in contexts near or adjacent to other domains and it may be possible to learn grammar-like rules which govern this.

It is worth noting that increasing the number of neighbours (increasing K) which functional roles can be inherited from degrades performance in this function-annotation task. Domains are typically involved in a large number of possible different protein functions. By increasing the number of neighbours GO terms can be inherited from the number of false positives is greatly increased and so performance degrades.

### Per Ontology Results

MCC values were also calculated for each of the three GO Ontologies (see table 3). Of the 2,358 GO terms used to annotate Eukaryotic sequences in Interpro: 1,018 are from the Molecular Function Ontology, 1,026 are from the Biological Process Ontology and 314 from the Cellular Component Ontology. The MCC values indicate different functional inheritance performance for each ontology with, unusually, the cellular component ontology being the best predicted set of terms. In the context of the vector embedding this may imply that the simple syntax contained in the domain orderings contains some additional information about where a protein is located within the cell. Of course these figures may also just reflect the degrees to which the prediction classes (True Positive, True Negative, etc…) are balanced for each of the ontologies.

**Table 3:** MCC values for K=1 nearest neighbour inheritance of GO terms, calculated for each separate GO ontology.

| Ontology | MCC |
| --- | --- |
| Biological Process | 0.27 |
| Molecular Function | 0.30 |
| Cellular Component | 0.34 |

In general we suspect the MCC calculated may underestimate the quality of the domain embedding. Using GO assignment to genes to annotate domains is inherently noisy. GO annotations may not be good descriptors of the specific role a domain plays within a given protein. Within the context of a multidomain proteins domains provide specific sub functionality such as providing a catalytic site, presenting one or more small molecule binding sites, providing membrane anchoring. It seems plausible if domains were annotated at a finer grained level, that better reflected these more specific roles, then the nearest neighbour assignment would return better results. The lack of a computer readable “domain ontology” remains an barrier for large scale studies of domain functionality and evolution.

### Vector mathematics on the domain embedding

One observation of semantic embeddings of natural languages is that arithmetic operations on the vectors have semantic or lexical meanings, one classic example being King – Man + Woman = Queen. We wished to investigate if simple vector arithmetic or translations for the protein domain embedding might have similar lexical meaning.

In the King to Queen example (see figure 4), subtracting Man from King takes you to a space in the embedding with the meaning of man “removed” such that adding the Woman vector will take you to Queen. We can perform similar vector subtractions for the domain embedding. In this context we would treat a domain’s set of GO terms as equivalent to its “meaning”, although, as discussed, this is a very lossy way to conceptualise the meaning of a domain. Nevertheless if we subtract two domain vectors we would hope the third vector is in a space where the remaining set of GO terms is the set difference of the two domains.

**Figure 4:**
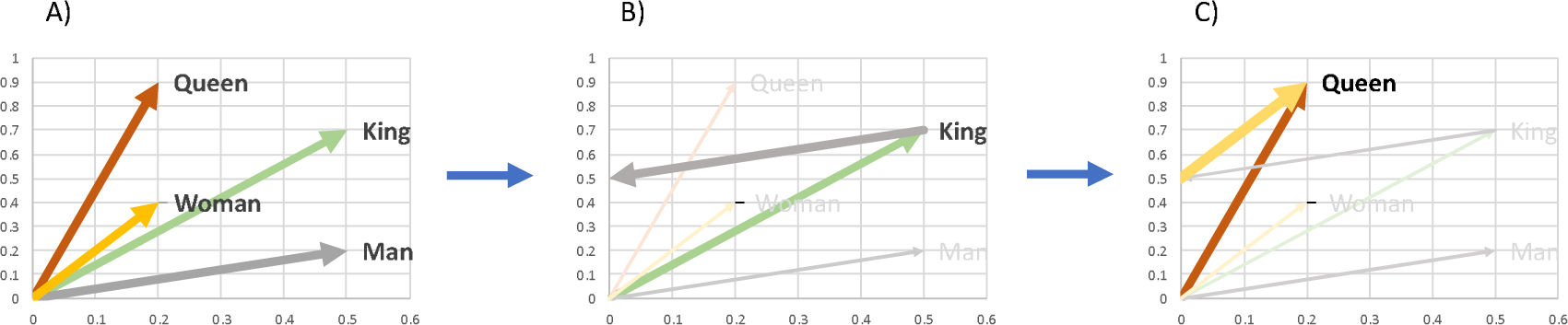
Example demonstrating semantically meaningful vector algebra. In A) four terms are placed in the vector space. If we subtract the **Man** vector from **King** (graph B), we move to an undefined point in the vector space. Adding the **Woman** vector (C) moves to the **Queen** vector.

We took the most common 20 Pfam domains, removing the one that isn’t present in eukaryotes and in turn subtracted all possible domain vectors. For the resulting third vector we found the nearest domain and tested the GO term overlaps with the initial two domains. In nearly all cases the resulting domain has minimal GO term overlaps with its parents. It is clear that this operation moves us to a region in the vector space where the domains’ “meaning” is profoundly altered, much as removing Man from King might be thought of as moving to a gender neutral space. What is not clear is what is the functional meaning of this in protein domain terms.

To investigate whether we could find more meaningful movements in the vector space we looked instead for translations in the vector space between mutually exclusive binary annotations. King and Queen are typically used as mutually exclusive labels that straddle some conceptual binary assignment (i.e. gender) and much the same is true of many GO terms. For instance in the Cellular Component Ontology annotation terms such as Intracellular and Extracellular might be viewed as a similar mutually exclusive binary.

We chose three binary cellular component term pairs; Intracellular (GO:0005622) vs Extracellular (GO:0005615), Nucleus (GO: 0005634) vs Cytoplasm (GO: 0005737) and Cytoplasm (GO: 0005737) vs transmembrane (GO: 0009279). For each pairing we identified proteins with domains annotated exclusively with one term and not the other term. Then for the first term we calculated the vector which moves from the location of the domain with the first term to the closest domain annotated with the second term. As with the prior analysis not having a detailed domain ontology prevents us from knowing if this closest domain is the most appropriate domain to move to. This led to a population of translation vectors which we could test to measure if the translation from a domain with one term to a domain with the other term was always vector oriented in a similar direction. We compared all Intracellular to Extracellular vectors in an all against all fashion and did the same for the other two pairs of terms (see Figure 5). If the translation is persevered in the vector space we would expect that all the vectors to have a small angle of deflection between them. In the transmembrane case there was no such alignment and not trend in the angles between the vectors. In both the Intracellular to Extracellular and the Nucleus to Cytoplasmic cases there is a clear distribution which peaks around 1.5 radians, indicating that in general the translation is commonly orthogonal and isn’t preserved in the vector space. However the intracellular to extracellular histogram shows a small leading tail below 1 radian (see figure 6) indicative of a small population of vectors which do approach alignment. And indeed we are able to find small numbers of genes in Interpro which share Pfam domains and where the difference is a substitution of an intracellular annotated domain for an extracellular domain such as G3I6X9 and A0A0L6WZ71 or I3L0A0 and G7Y5H3.

**Figure 5:**
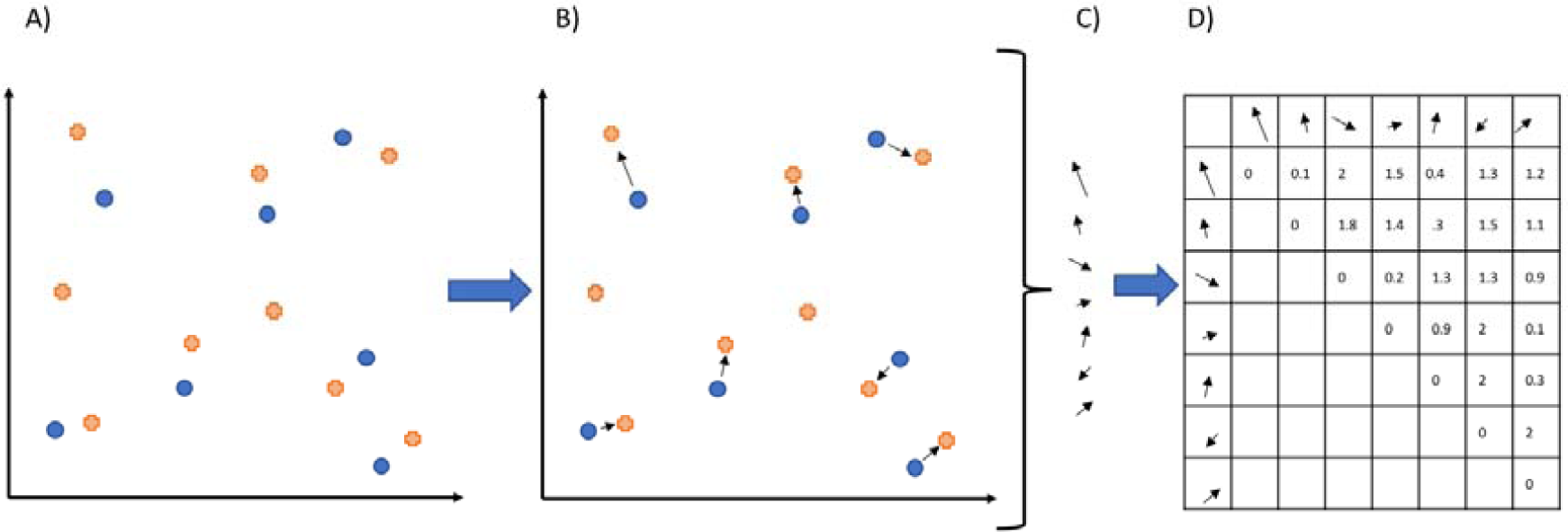
Comparing translation vector from one binary GO property to another. A) Putative vector embedding of Intracellular (blue dots) and Extracellular (orange crosses) labelled domains. B) Vectors which translate each intracellular domain to its closest Extracellular labelled domain. C) Vectors are extracted and pooled D) Angle between each vector is compared to find vectors that point in the same direction.

**Figure 6:**
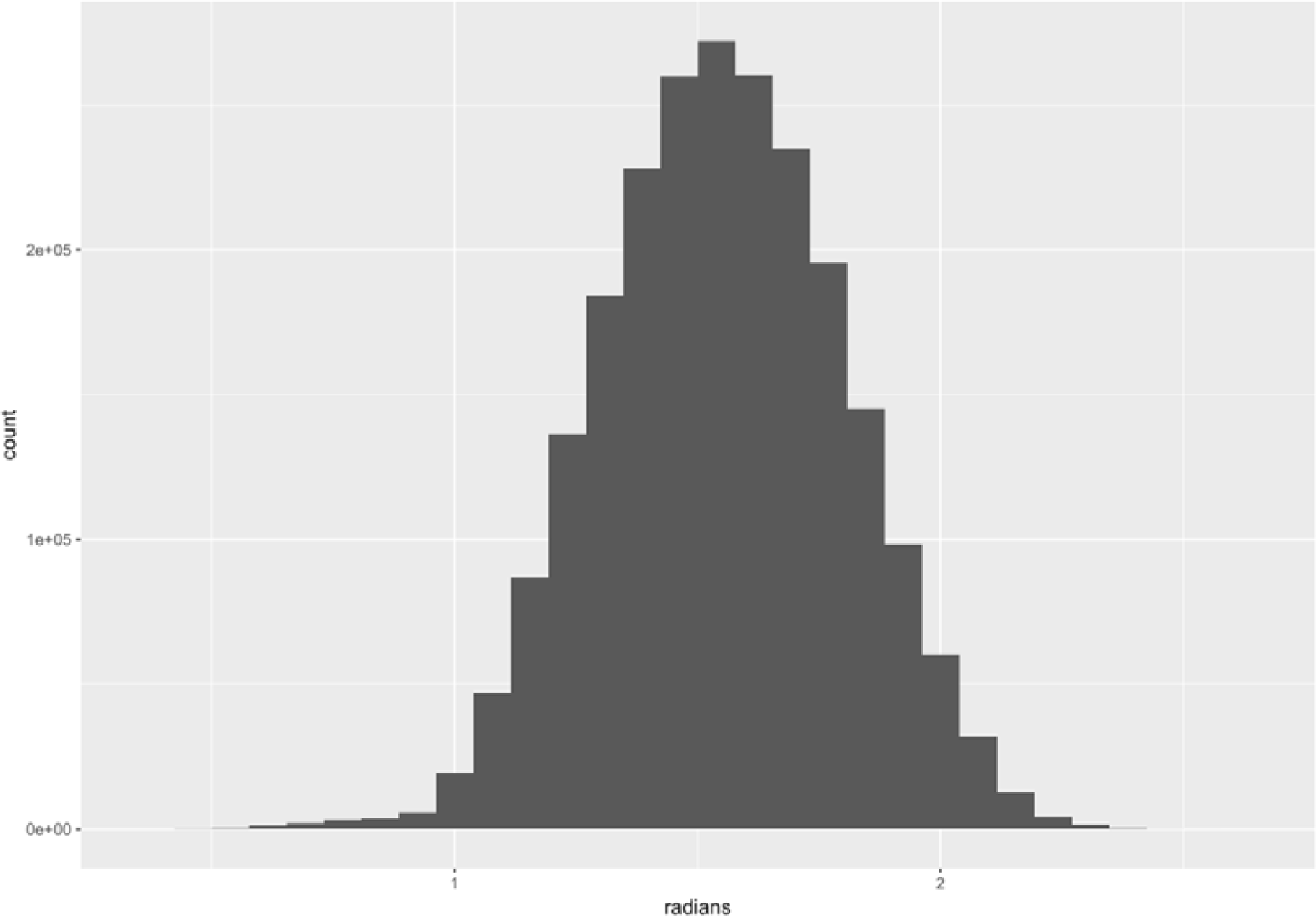
Histogram of transformation vector angles. For intracellular to extracellular.

### Domains of Unknown Function

As the word2vec embedding has some meaningful structure with regards GO term inheritance we can also use a nearest neighbour approach to suggest putative sets of GO terms that each eukaryotic Pfam Domain of Unknown Function (Pfam DUFs) may take part in. Our corpus of eukaryotic genes contained annotations from 3,918 DUFs. Using a single nearest neighbour inheritance method 1,292 of these domains could be assigned new GO terms (i.e. their nearest neighbour in the embedding was annotated and was not a gap or other sequence region). On average each DUF gets 11 novel GO terms assigned. In figure 7 the distribution of terms indicates that the majority of DUFs receive only a handful of putative GO assignments. We suggest that such assignments could be used as starting points for Pfam domain annotations and with relatively fewer terms to confirm in most these shouldn’t make such annotation tasks more onerous or obfuscated. We make these annotations available (see supplementary material) and note they could make a starting point for future annotation of these domain in Pfam.

**Figure 7:**
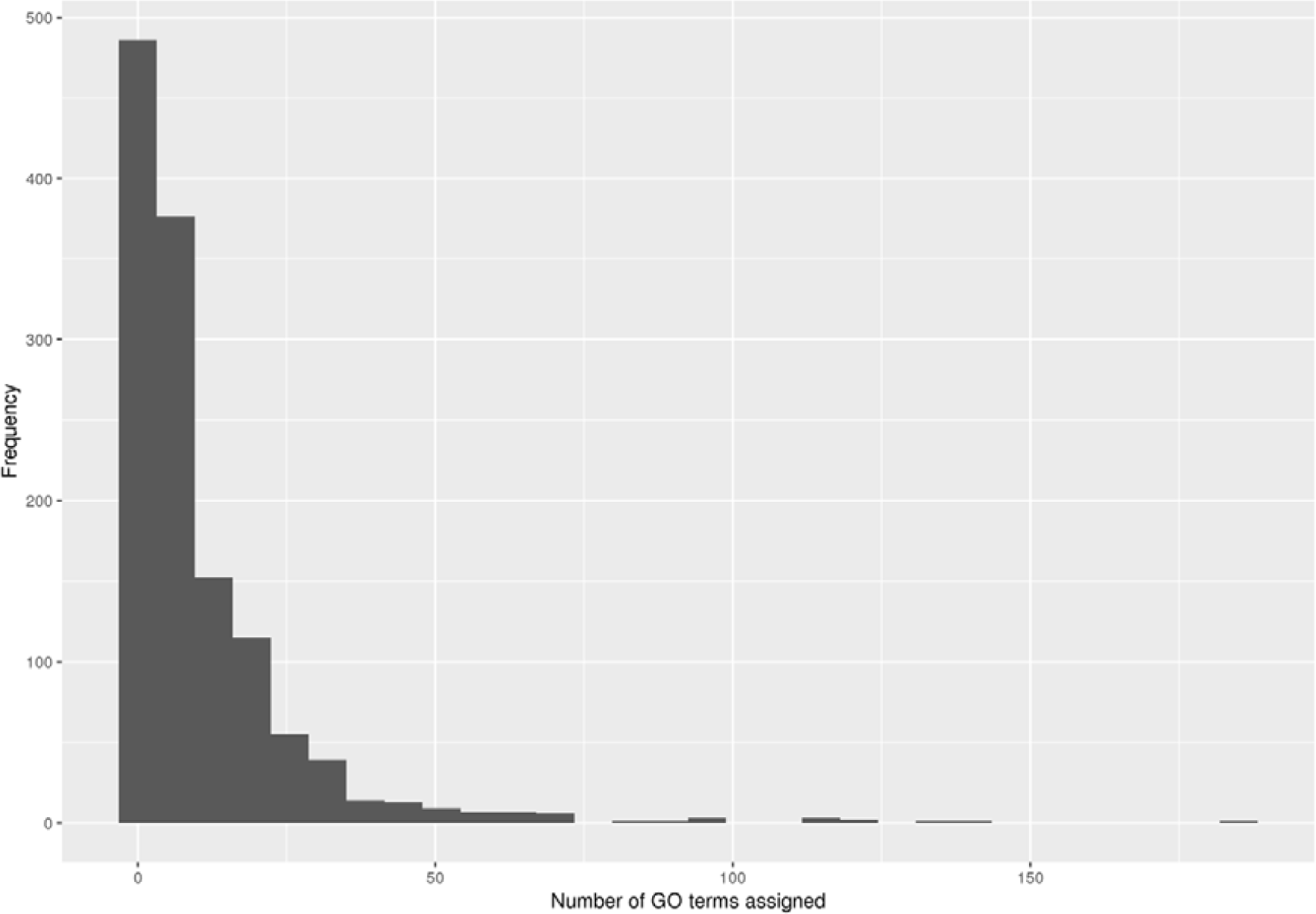
Frequency of the number of GO terms assigned to DUFs Discussion

## Discussion

Applying word2vec to protein domains, making the assumption that multi-domain proteins are sentence-like, reveals that domains display some manner of semantic or lexical structure. Given this it should be possible in future to elucidate statistical rules for domain placement in multi-domain proteins which in turn would have applications in protein design and modelling. Though whatever statistical propensities for domain placement that may exist may be commonly or trivially ignored by evolution.

Additionally word2vec was trained and designed to work over very large corpuses of human language. The nine million Eukatyotic Interpro sequences used in this study may represent too small a corpus of “sentences” to develop a higher quality embedding of word-tokens. Additionally multi-domain proteins typically have fewer than six domains whereas human sentences are frequently longer. This means sequential sets of domains are unlikely to be sufficiently analogous sentences which may also hinder the performance of the vector embedding. All these issue may be address by retraining or developing a new word2vec-like method better optimised for domain embeddings of sets of protein domains.

Using GO annotations to annotate domains is necessarily noisy. It is not clear that they are the best way to encode the lexical “meaning” of a domain in its multi-domain context. In future a finer grained annotation of domains’ sub-functional roles will be necessary to correctly interpret the lexical meaning of arithmetic transformations of vectors in the embedding space. Nevertheless this work does open up the tantalising possibility that protein domains have contextual lexical meaning that could be used to derive rules for multidomain protein evolution.

### Code & Data

All code is available on github and the domain assignments, genism model, token distance matrix and DUF assignements are available via our webserver https://github.com/psipred/domain_word2vec_scripts https://bioinfadmin.cs.ucl.ac.uk/downloads/word2vec/

